# Nucleus accumbens neurochemistry in human anxiety: A 7 T ^1^H-MRS study

**DOI:** 10.1101/498337

**Authors:** Alina Strasser, Lijing Xin, Rolf Gruetter, Carmen Sandi

## Abstract

Individual differences in anxiety provide a differential predisposition to develop neuropsychiatric disorders. The neurochemical underpinnings of anxiety remain elusive, particularly in deep structures, such as the nucleus accumbens (NAc) whose involvement in anxiety is being increasingly recognized. We examined the associations between the neurochemical profile of human NAc metabolites involved in neural excitation and inhibition and inter-individual variation in temperamental and situational anxiety. Twenty-seven healthy 20-30 years-old human males were phenotyped with questionnaires for state and trait anxiety (State-Trait Anxiety Inventory, STAI), social anxiety (Liebowitz Social Anxiety Scale), depression (Beck Depression Inventory, BDI) and fatigue (Mental and Physical State Energy and Fatigue Scales, SEF). Using proton magnetic resonance spectroscopy (^1^H-MRS) at 7 Tesla (7T), we measured metabolite levels for glutamate, glutamine, GABA and taurine in the NAc with. Salivary cortisol was also measured. Strikingly, trait anxiety was negatively associated with NAc taurine content. Perceived situational stress was negatively associated with NAc GABA, while positively with the Glu/GABA ratio. These findings were specific, as no correlation was observed between NAc taurine or GABA and other phenotypic variables examined (i.e., state anxiety, social anxiety, depression, or cortisol), except for a negative correlation between taurine and state physical fatigue. This first 7T study of NAc neurochemistry shows relevant metabolite associations with individual variation in anxiety traits and situational stress and state anxiety measurements. The novel identified association between NAc taurine levels and trait anxiety may pave the way for clinical studies aimed at identifying new treatments for anxiety and related disorders.

## Introduction

Anxiety is an emotion characterized by apprehension or dread and enhanced vigilance, frequently accompanied by behavioral, cognitive and physiological stress responses. Although anxiety can help the individual preparing to detect and deal with threats (Bateson et al., 2011), when excessive, it can be maladaptive or pathological and manifest in a variety of disorders (Price, 2003).

Individuals differ greatly in their behavioral and physiological responses to actual or potential threats, following a continuum from mild to strong. While ‘state’ anxiety refers to transitory emotional and motivational state displayed in response to immediate uncertainty (Cattell, 1966; Eysenck et al., 2007; Spielberger, 1972), ‘trait’ anxiety is the stable predisposition of an individual to judge environmental events as potentially threatening (Domschke and Reif, 2012; Mann et al., 2012; Sih et al., 2004; Toye and Cox, 2001). High trait anxious individuals are at risk to develop stress-induced neuropsychiatric disorders, particularly anxiety disorders and depression (Rogers et al., 2013; Sandi and Richter-Levin, 2009; Weger and Sandi, 2018). ‘Social’ anxiety refers to persistent fear in social interactions and performance (Heimberg et al., 1999) that, at high levels, becomes social anxiety disorder (formerly known as social phobia) (Heimberg et al., 1999).

Identifying neurobiological factors related to individual differences in temperamental and situational anxiety will advance our understanding of mechanisms underlying stress vulnerability and might lead to the development of new treatments for anxiety disorders. However, the neurobiology of anxiety is scarcely understood (Craske et al., 2017; Weger and Sandi, 2018). Previous work has highlighted imbalances in neural excitation and inhibition (E/I) in several brain areas in anxiety and mood disorders) and following stress exposure (Cordero et al., 2016; Hasler et al., 2010; Houtepen et al., 2017; Moghaddam, 2002; Popoli et al., 2012; Sandi, 2011). In addition, taurine, an abundant free β-amino acid, has been shown to act as endogenous inhibitor of cellular excitability and network activity (Davison and Kaczmarek, 1971; El Idrissi and Trenkner, 2004; Jia et al., 2008). Notably, taurine treatment is effective in reducing anxiety-like behaviors (Mezzomo et al., 2016; Zhang and Kim, 2007). However, to date, there is no information about taurine levels in the human brain in the context of anxiety.

Recently, a role for the nucleus accumbens (NAc), the main anatomical constituent of the ventral striatum (VS), is emerging in the context of anxiety (Gunaydin and Kreitzer, 2016; Levita et al., 2012) interconnected with the NAc function in behavioral adaptation (Haber and Behrens, 2014). In rodents, anxiety-like behaviors can be regulated by pharmacological (Heshmati et al., 2016; Lopes et al., 2012) or genetic (Crofton et al., 2017; Feng et al., 2017; Shen et al., 2016; Zhao and Gammie, 2018) manipulation of NAc neurochemistry. Importantly, anxiety-like behaviors have also been related to variations in NAc mitochondrial function (Hollis et al., 2015; Van Der Kooij et al., 2018) and brain energy metabolism (Larrieu et al., 2017). A few studies have also highlighted NAc structural (Kühn et al., 2011) and functional (Goff et al., 2013; Levita et al., 2012) alterations in association with anxiety and depression in humans. However, there is virtually no information on the NAc/VS neurochemical profile, nor on its relationship with anxiety.

Therefore, the aim of this study was to perform proton magnetic resonance spectroscopy (^1^H-MRS) at 7 Tesla (7 T) in the NAc of individuals phenotyped for anxiety (state, trait and social) and self-perceived situational stress, with a focus on glutamate, glutamine, GABA and taurine.

## Experimental Procedures

### Participants

Thirty-eight healthy males, 20-30 years old, from the University of Lausanne and the Ecole Polytechnique Fédérale de Lausanne (EPFL) were recruited for the study. All participants gave informed consent before participating and were debriefed after the experiment. The Cantonal Ethics Committee of Vaud, Switzerland, approved the study protocol. Phenotypic assessments and MRS measurements for neurochemical assessment were performed on two different days for the two brain regions, i.e. nucleus accumbens (NAc; *N* = 27) and occipital lobe (OL; *N* = 17).

Inclusion criteria for the study were being male, 20-30 years old, right-handed, no regular drug or medication intake, non-smoker, no history of psychiatric or neurological illness, and no metallic implants. Participants were instructed not to drink any caffeinated drinks one hour before the experiment, and not to be hungry or thirsty when arriving in the laboratory. Furthermore, they were instructed not to take any medication and to avoid physical effort within 24 hours before the experiment.

### Psychometric questionnaires

State and trait anxiety were assessed with the State-Trait Anxiety Inventory (STAI) (Spielberger, 2010). Self-perceived stress was measured with a 10-point rating scale on the scanner bed 30 min before ^1^H-MRS. Social anxiety and depression were assessed with the Liebowitz Social Anxiety Scale (LSAS) (Heimberg et al. 1999) and the Beck Depression Inventory (BDI) (Beck, 1961). Other questionnaires included the Mental and Physical State Energy and Fatigue Scales (SEF) (Loy and O’Connor, 2016). Tables S1 and S2 show the phenotypic characteristics of participants for the questionnaires filled in each of the two experimental sessions. For further information, see Supplementary Materials.

### Salivary cortisol

Saliva was collected three times (baseline, pre and post MRS acquisition) on the first experimental day. Free cortisol concentrations were determined with the Salimetrics Salivary Cortisol Enzyme Immunoassay Kit. Area under the curve values (AUC_G_, AUC_I_) were computed according to Pruessner et al. (2003) (Pruessner et al., 2003). Table S3 includes the descriptive statistics for the salivary cortisol concentrations. For further information, see Supplementary Materials.

### ^1^H magnetic resonance spectroscopy (^1^H-MRS)

^1^H-MRS was performed on a 7 T/68 cm MR scanner (Magnetom, Siemens Medical Solutions, Erlangen, Germany) equipped with head gradients with maximum strengths of 50 mT/m and 2^nd^-order shims of up to 13.5 mT/m^2^ for Z^2^ and 11.8 mT/m^2^ for ZX, ZY, X^2^-Y^2^ and XY. A single-channel quadrature transmit and a 32-channel receive coil (Nova Medical Inc., MA, USA) were used for ^1^H-MRS in the NAc. A home-built ^1^H quadrature transmit/receive surface coil was used for OL ^1^H-MRS.

To ensure the full inclusion of the NAc, the VOI was defined by the third ventricle medially, the subcallosal area inferiorly, and the body of the caudate nucleus and the putamen laterally and superiorly, in line with definitions of NAc anatomy identifiable on MRIs (Neto et al., 2008) (Figure 1A). Localized single-voxel ^1^H-MR spectra from the NAc were obtained from twenty-seven participants. ^1^H-MRS in the OL was performed as the experimental control (*N* = 17). For detailed information on ^1^H-MRS procedures, see Supplementary Materials.

**Figure 1.**
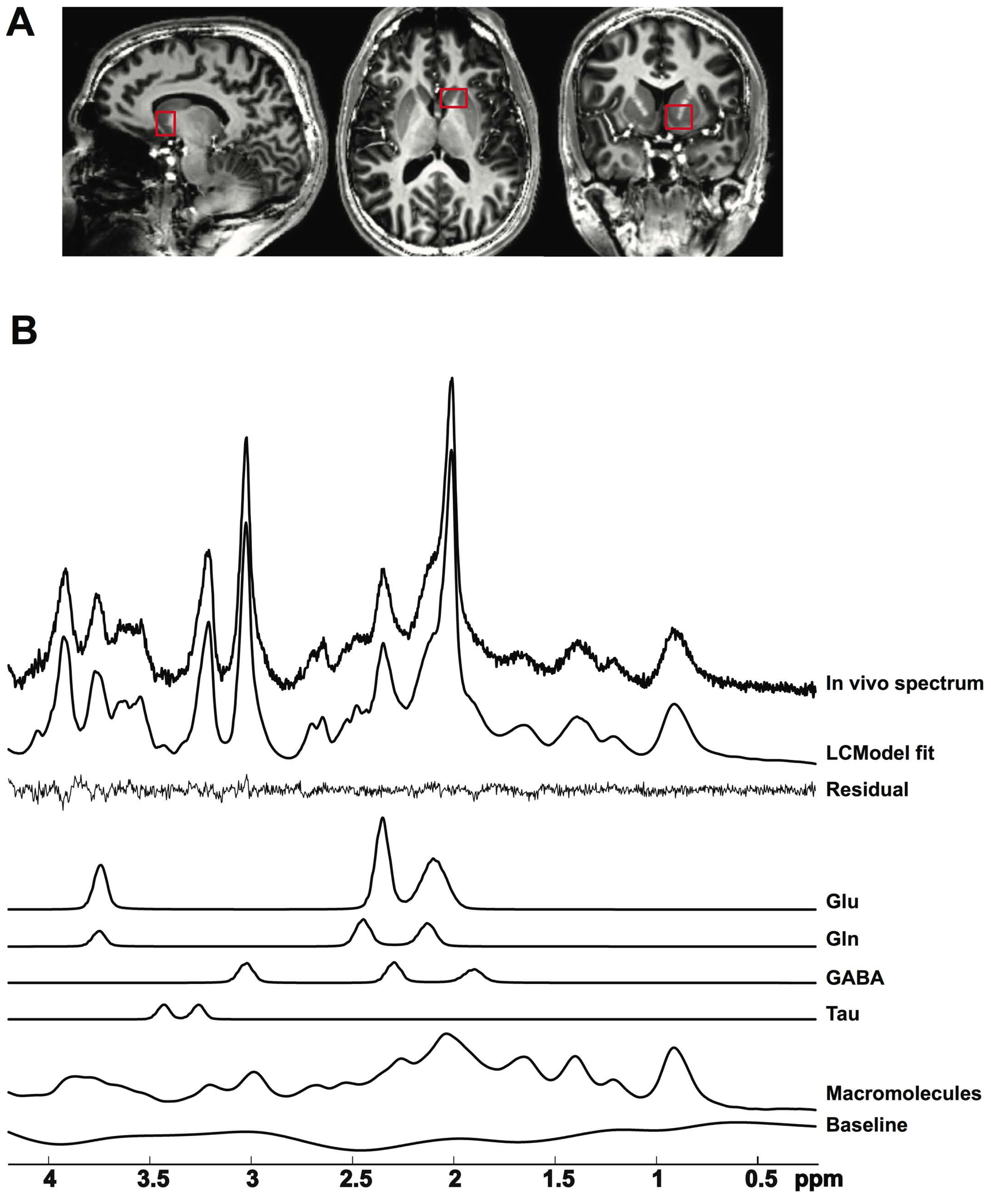
In vivo ^1^H-MR spectroscopy in the nucleus accumbens. (**A**) MP2RAGE images with the volume of interest (14 × 10 × 13 mm^3^) in the left nucleus accumbens for localized ^1^H-MRS. (**B**) Representative ^1^H-MR spectrum acquired with the semi-adiabatic SPECIAL sequence at 7 T (TE/TR = 16 ms/6500 ms, 256 averages), the corresponding LCModel spectral fit, fit residual, macromolecules, baseline and individual metabolite fits for glutamate (Glu), glutamine (Gln), gamma-aminobutyric acid (GABA) and taurine (Tau).

### ^1^H-MRS data processing

1H-MR spectra were corrected, summed and analyzed using LCModel with a basis set including simulated metabolite spectra and an experimentally measured macromolecule baseline. The spectral analysis range was set between 0.2 and 4.2 ppm (Govindaraju et al., 2000; Provencher, 2001; Schaller et al., 2014).

MP2RAGE images were segmented for VOI tissue decomposition into grey matter (GM), white matter (WM) and cerebrospinal fluid (CSF). Water concentrations within VOIs were calculated based on the segmentation values assuming water concentrations of 43300 mM in GM, 35880 mM in WM, and 55556 mM in CSF. Metabolite concentrations were then partial-volume corrected for the CSF fraction.

The signal-to-noise ratio (SNR) was obtained using the N-acetylaspartate (NAA) peak height at 2.01 ppm divided by the SD of the noise between 9.5 - 10 ppm. FWHM (ppm) was given by the output of LCModel.

Metabolite concentrations with relative Cramér-Rao lower bounds (CRLB) of 999 % were considered as non-detectable. No threshold of relative CRLB (%) was used. Absolute CRLBs were calculated by multiplying metabolite concentration with relative CRLB (%). Total glutamate and glutamine are reported as Glx. Metabolite concentrations are reported in μmol/g. Excitation/inhibition (E/I) balance is reported as Glu/GABA and Glx/GABA.

### Statistical analyses

Data was processed and analyzed in Microsoft Excel 2011, IBM SPSS Statistics 20, Cocor, MATLAB R2017a, and GraphPad Prism 7. Normality testing was done with the Shapiro-Wilk test with statistical significance at *P* < 0.05. Associations were quantified with the Pearson’s correlation coefficient for normally distributed variable pairs. Associations including a not normally distributed variable were quantified with Spearman’s rank correlation coefficient. All statistical significance testing was performed two-tailed. *P*-values. Unless stated in the text, *P*-values were not adjusted for multiple comparisons.

To test the specificity of NAc metabolite associations with anxiety phenotypes, the differences in the observed correlations in the NAc versus the OL were tested for statistical significance with a modification of Dunn and Clark’s *Z* using Fisher’s *Z* procedure and Zou’s confidence interval (CI) (Zou, 2007), applying the OL ^1^H-MRS sample size (*N* = 17).

## Results

### Localized single-voxel ^1^H-MR spectra

NAc VOI tissue fractions of 85 ± 4 % GM, 12 ± 5 % WM, and 2 ± 3 % CSF were obtained by MP2RAGE image segmentation. A typical in vivo ^1^H-MR spectrum of the NAc and its LCModel fit are shown in Figure 1B. Twenty-eight min acquisition of ^1^H-MR spectra in the NAc (VOI = 1.82 ml) resulted in a spectral SNR of 72 ± 9. Optimizing 1^st^- and 2^nd^-order shims by FAST(EST)MAP resulted in FWHM of 0.048 ± 0.006 ppm. Concentrations and CRLBs of the NAc metabolites of interest are reported in Table S4, and absolute CRLBs for glutamate (Glu), glutamine (Gln), GABA and taurine were in a similar range. In the OL (VOI = 10 ml), spectra were obtained at a spectral SNR of 237 ± 51, FWHM of 0.031 ± 0.004 ppm, and with 62 ± 5 % GM, 19 ± 4 % WM, and 19 ± 4 % CSF. NAc metabolite inter-correlations are shown in Table S5. For the OL, the corresponding information and associations are shown in Tables S6 and S7.

### Associations between anxiety phenotypes and situational stress with nucleus accumbens neurochemistry

Figure 2A shows the correlation matrix for the different anxiety phenotypes (state, trait and social anxiety and self-perceived situational stress) and NAc metabolites (Glu, Gln, Glx, GABA and taurine). We found trait anxiety to be negatively correlated with NAc taurine (Figure 2A, 2B and S2). Importantly, this correlation remained significant after Bonferroni correction for five comparisons (α = 0.05/5 = 0.01). No significant associations were found between trait anxiety and any of the other metabolites (i.e., Glu, Gln, Glx, and GABA), nor between any of the analyzed metabolites and state anxiety or social anxiety (Figure 2A). Age was not associated with metabolite concentrations or E/I balance in the NAc (Figure S3) and was not added as a covariate in the statistical analyses. Furthermore, in the OL, we found no evidence for significant associations between the analyzed metabolites and state anxiety, trait anxiety, or situational stress (Figure S4). However, given that the sample size for the OL control experiment was lower than for the NAc experiment, we performed further analyses to compare the slopes of the two correlations (i.e., for NAc and OL). The correlation between trait anxiety NAc taurine was significantly different to the one between trait anxiety and OL taurine (Figure S4A; *Z* = −1.926, *P* = 0.05), as confirmed by Zou’s test (95% CI did not include 0; − 1.148 to −0.003).

**Figure 2.**
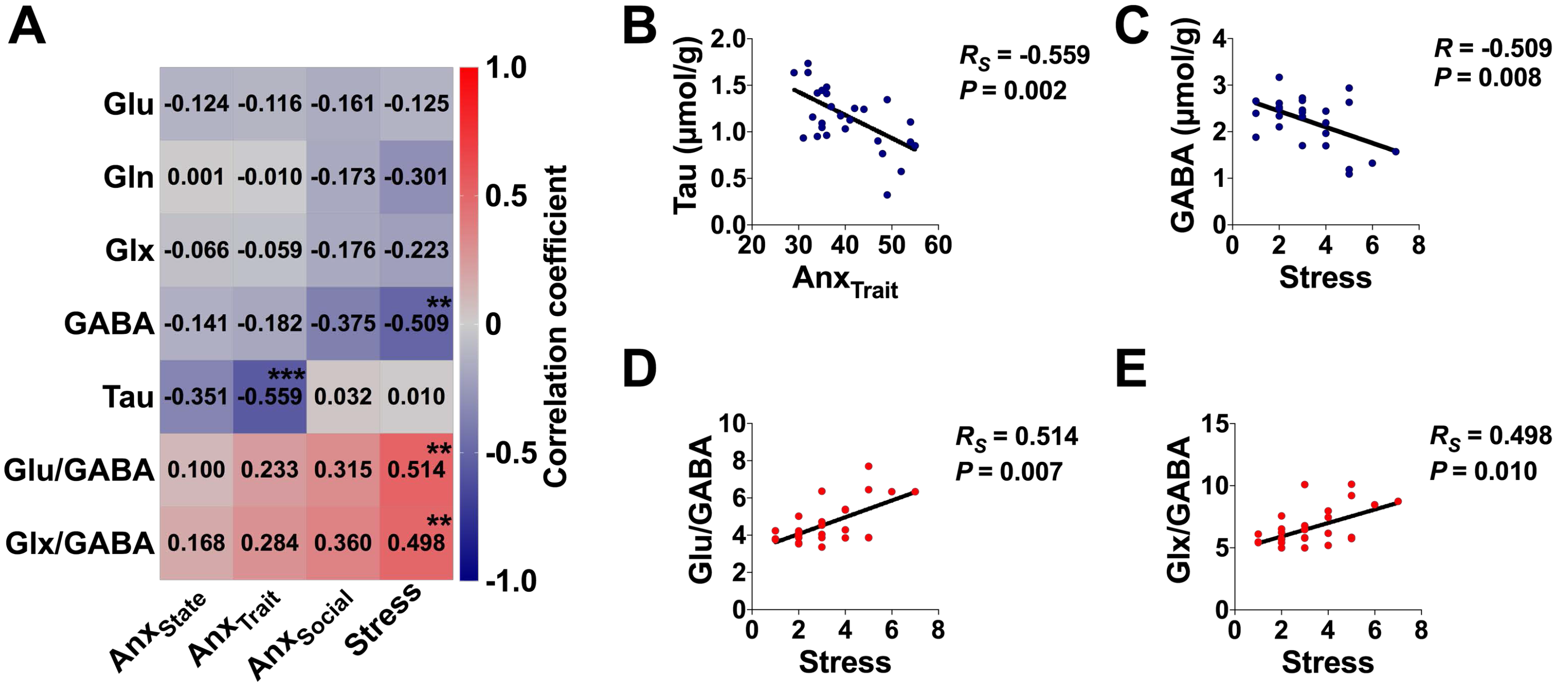
Associations between anxiety phenotypes and acute stress with neurochemistry in the nucleus accumbens. (**A**) Correlation matrix between state anxiety, trait anxiety, social anxiety, acute stress, depression and metabolites and E/I balance. Each cell contains the correlation coefficient corresponding to the color scale. (**B**) Scatter plot of trait anxiety and taurine. (**C**) Scatter plot of acute stress and GABA. (**D**) Scatter plot of acute stress and Glu/GABA. (**E**) Scatter plot of acute stress and Glx/GABA. Anx_State_, state anxiety (STAI-S), *N* = 27; Anx_Trait_, trait anxiety (STAI-T), *N* = 27; Anx_Social_, social anxiety (LSAS), *N* = 26; Stress, self-perceived acute stress, *N* = 26. Glu, glutamate; Gln, glutamine; Glx, Glu+Gln; GABA, gamma-aminobutyric acid; Tau, taurine. *R*, Pearson’s correlation coefficient. *R_S_* = Spearman’s correlation coefficient. \*\**P* < 0.01; \*\*\**P* < 0.001.

In addition, we found a significant negative correlation between situational stress experienced during the preparation for the scan (30 min prior to spectra acquisition) and GABA in the NAc (Figure 2A and 2C). This correlation remained significant after Bonferroni correction for five comparisons (α = 0.05/5 = 0.01). In the OL, there was a trend towards significance in the correlation between situational stress and GABA (Figure S4B) and the correlations in the two brain regions were not significantly different from each other (*Z* = −0.594, *P* = 0.553), as also confirmed by the Zou test (95% CI included 0; −0.857 to 0.472).

### Associations between anxiety phenotypes and situational stress with nucleus accumbens E/I metabolite ratios

Figure 2A shows the correlation matrix between the different anxiety phenotypes (state, trait and social anxiety and situational stress) and NAc E/I metabolite ratios (Glu/GABA and Glx/GABA). Only situational stress showed significant correlations: it was positively correlated with both, NAc Glu/GABA (Figure 2D) and Glx/GABA (Figure 2E). The association between situational stress and Glu/GABA survived Bonferroni correction for five comparisons (α = 0.05/5 = 0.01). None of these analyses reached significance when performed on OL metabolites (Figure 5S). However, the correlations between situational stress and either NAc Glu/GABA [(*Z* = 1.184, *P* = 0.237); as confirmed by the Zou test (95% CI included 0; −0.273 to 1.021)] or NAc Glx/GABA [(*Z* = 1.155, *P* = 0.248); as confirmed by the Zou test (95% CI included 0; −0.293 to 1.003)] were not significantly different from the corresponding ones performed in the OL.

### Associations between anxiety trait and situational stress with depression, energy and fatigue

Our previous analyses showed that trait anxiety correlates negatively with NAc taurine and situational stress correlates negatively with GABA (Figure 2). Here, we asked whether trait anxiety and situational stress scores are related to depression and to mental and physical energy or fatigue. Figure 3A shows the corresponding correlation matrix. Trait anxiety was positively correlated with depression (Figure 3B). We also found that trait anxiety was positively correlated with state and trait mental fatigue (Figure 3C and Figure 3D) as well as with state and trait physical fatigue (Figure 3E and Figure 3F).

**Figure 3.**
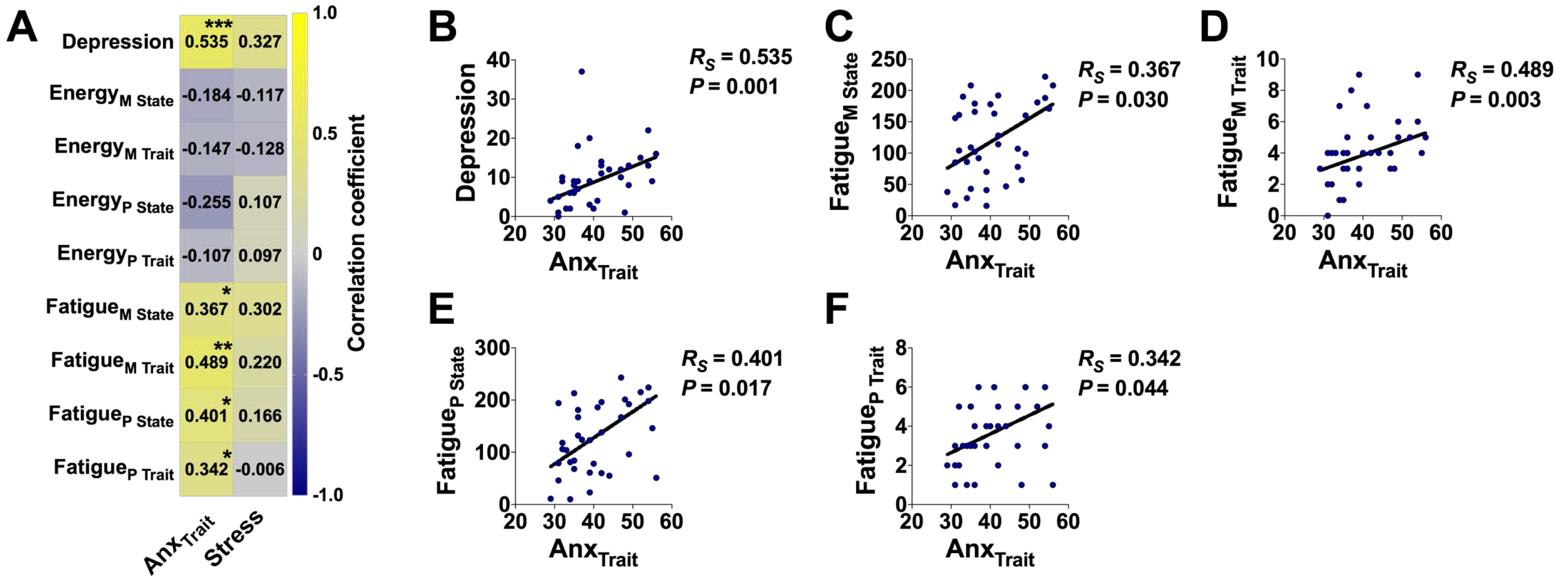
Associations between anxiety, acute stress, depression and mental and physical energy and fatigue. (**A**) Correlation matrix of trait anxiety and acute stress with depression and mental and physical energy and fatigue. Each cell contains the correlation coefficient corresponding to the color scale. (**B**) Scatter plot of trait anxiety and depression. (**C**) Scatter plot of trait anxiety and state mental fatigue. (**D**) Scatter plot of trait anxiety and trait mental fatigue. (**E**) Scatter plot of trait anxiety and state physical fatigue. (**F**) Scatter plot of trait anxiety and trait physical fatigue. Anx_Trait_, trait anxiety (STAI-T), *N* = 35; Stress, self-perceived acute stress, *N* = 35; Depression (BDI), *N* = 35; Energy_M State_, state mental energy; Energy_M Trait_, trait mental energy; Energy_P State_, state physical energy; Energy_P Trait_, trait physical energy; Fatigue_M State_, state mental fatigue; Fatigue_M Trait_, trait mental fatigue; Fatigue_P State_, state physical fatigue; Fatigue_P Trait_, trait physical fatigue. *R_S_* = Spearman’s correlation coefficient. \**P* < 0.05; \*\**P* < 0.01; \*\*\**P* < 0.001.

### Associations of depression and fatigue with nucleus accumbens neurochemistry

Therefore, given the findings of associations between trait anxiety and depression and the energy and fatigue phenotypes (reported in Figure 3), we further explored the potential associations between these parameters and NAc metabolites (see Figure 4A). State physical fatigue was negatively correlated with taurine (Figure 4B), and the correlation between state physical fatigue and NAc taurine was significantly different to the one between state physical fatigue and OL taurine (Figure S6; *Z* = − 2.129, *P* = 0.033), confirmed by the Zou test (95% CI did not include; −1.202 to −0.075). In addition, given that trait anxiety is a strong risk factor for the development of depression, and there is a large comorbidity between anxiety disorders and depression (Sandi and Richter-Levin, 2009), we further analyzed whether the correlation between NAc taurine and trait anxiety (Figure 2B) was significantly different to the one between NAc taurine and depression [note that depression did not significantly correlate with any of the analyzed metabolites, including taurine; Figure 4A; *Z* = −3.2541, *P* = 0.001). Zou’s test confirmed this finding (95% CI did not include 0; −0.906 to −0.244)].

**Figure 4.**
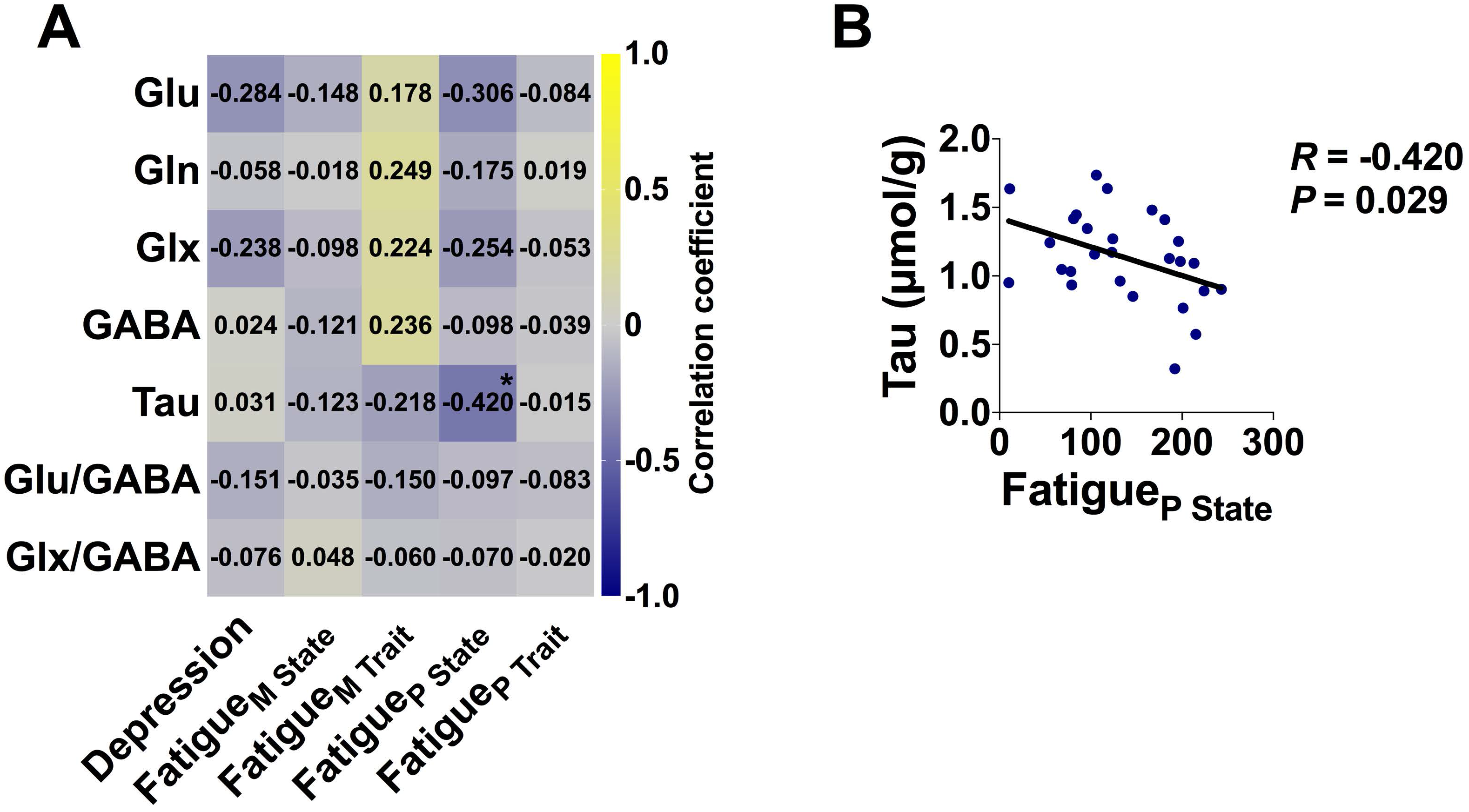
Associations between depression, mental and physical energy and fatigue and neurochemistry in the nucleus accumbens. (**A**) Correlation matrix between mental and physical fatigue and metabolites and E/I balance. Each cell contains the correlation coefficient corresponding to the color scale. (**B**) Scatter plot of state physical fatigue and taurine. Depression (BDI), *N* = 35; Fatigue_M State_, state mental fatigue; Fatigue_M Trait_, trait mental fatigue; Fatigue_P State_, state physical fatigue; Fatigue_P Trait_, trait physical fatigue. Glu, glutamate; Gln, glutamine; Glx, Glu+Gln; GABA, gamma-aminobutyric acid; Tau, taurine. *R*, Pearson’s correlation coefficient. \**P* < 0.05.

**Figure 5.**
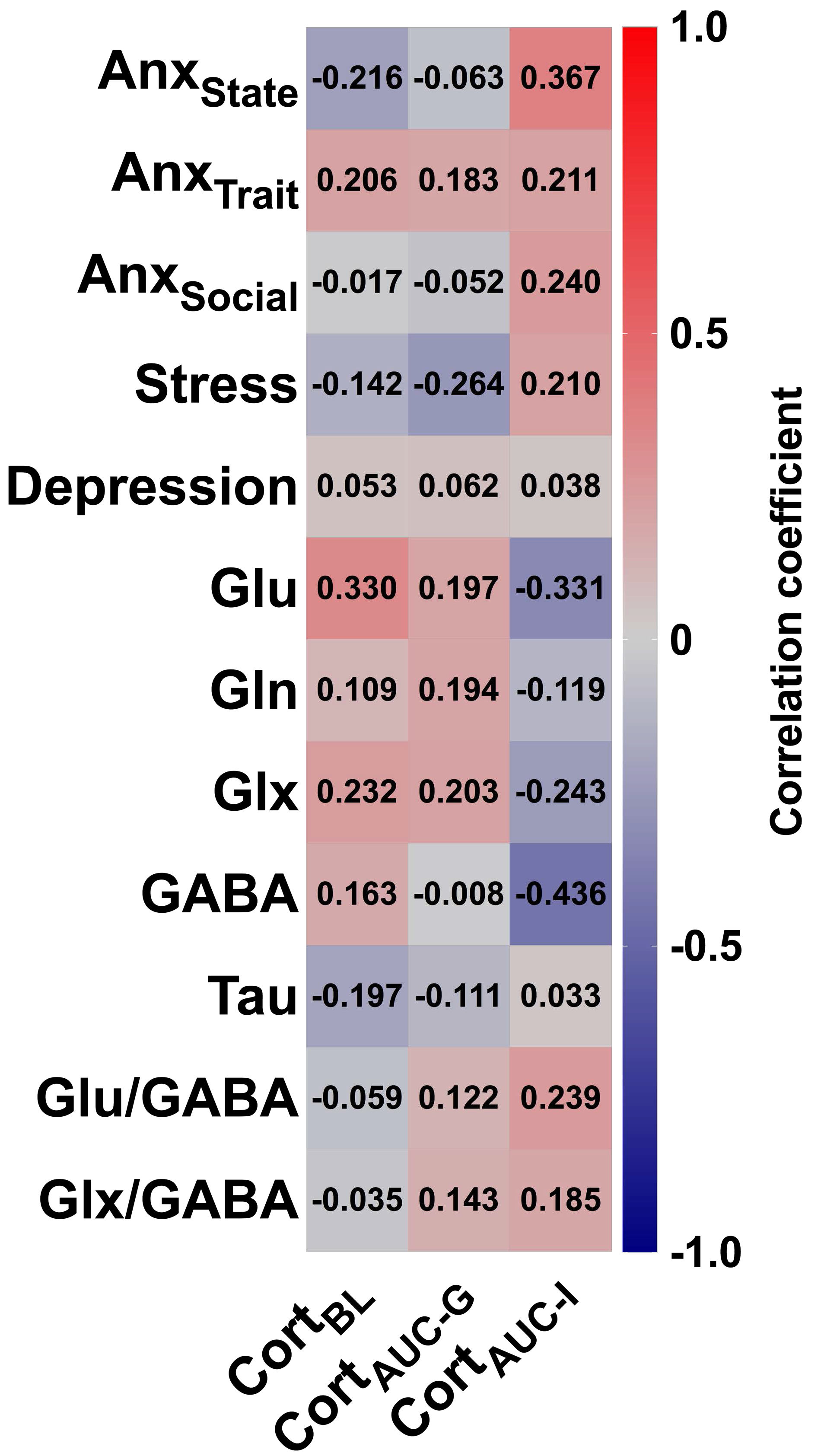
Correlation matrix between salivary cortisol and personality phenotypes and neurochemistry in the nucleus accumbens. Cort_BL_, cortisol baseline; Cort_AUC-G_, cortisol area under the curve with respect to ground; Cort_AUC-I_, cortisol area under the curve with respect to increase. Anx_State_, state anxiety (STAI-S); Anx_Trait_, trait anxiety (STAI-T); Anx_Social_, social anxiety (LSAS); Stress, self-reported stress. Glu, glutamate; Gln, glutamine; Glx, Glu+Gln; GABA, gamma-aminobutyric acid; Tau, taurine. Each cell contains the correlation coefficient corresponding to the color scale.

### Lack of association between salivary cortisol and personality variables or nucleus accumbens neurochemistry

Finally, we analyzed potential correlations between salivary cortisol levels, collected at different time points during participants’ exposure to the first experimental session, and the different anxiety and stress phenotypes, as well as with NAc metabolites. As shown in Figure 5, we found no evidence for significant correlations, with the exception of a tendency towards significance in the negative correlation between Cort_AUC-I_ [representing endocrine system sensitivity in terms of changes over time (Pruessner et al., 2003)] and NAc GABA levels (*R* = −0.436, *P* = 0.055, *N* = 20).

## Discussion

In this 7 T ^1^H-MRS study of the human NAc, we investigated the associations between anxiety phenotypes and NAc metabolites involved in neural excitation and inhibition, including Glu, Gln and GABA, as well as the β-amino sulfonic acid taurine. We report a marked positive correlation between trait anxiety and NAc taurine content, and a negative correlation between perceived situational stress and NAc GABA, while positive with the Glu/GABA ratio. These findings were specific, as no correlation was observed between NAc taurine or GABA and other phenotypic variables examined (i.e., state anxiety, social anxiety, depression, or cortisol), except for a negative correlation between taurine and state physical fatigue.

To the best of our knowledge, this study constitutes the first report of the neurochemical profile of the human NAc using ^1^H-MRS at 7 T. Performing ^1^H-MRS in the NAc is challenging because its size is relatively small (approx. 1.5 cm^3^) and because the large local susceptibility gradient results in broad spectral linewidths, which can further degrade signal intensity. We used a short-TE (16 ms) ^1^H-MRS technique together with 7 T to enhance chemical shift dispersion and increase measurement sensitivity. At lower magnetic field strength (< 7 T), the reliability of the separate measurement of Glu and Gln is more challenging due to the limited peak separation. However, applying 7 T, we were able to reliably separate the Glu and Gln peaks, which in our study is supported by the low correlation coefficient (from −0.12 to 0.07; *N* = 27). A recent VS ^1^H-MRS study at 3 T, could only measure a limited number of metabolites (Liu et al., 2017), due to the lower sensitivity and spectral dispersion available at 3 T as well as the longer TE used (40 ms). Furthermore, we did not use relative CRLB (%) as a threshold for reporting metabolite concentrations. It has been suggested that applying 20-50 % CRLB as a threshold for data exclusion might lead to a biased lack in reporting metabolites at low tissue concentration (Kreis, 2016).

Our finding linking NAc taurine concentrations with natural variations in trait anxiety in healthy humans seems remarkable, given the existence of studies in rodents (Chen et al., 2004; El Idrissi et al., 2009; Kong et al., 2006; Zhang and Kim, 2007) and zebrafish (Mezzomo et al., 2017, 2016) have reported reductions in anxiety-like behaviors following treatment with taurine. Information about taurine in the human brain in the context of anxiety had previously been lacking. Indirect evidence from studies in rodents supports our observed link between anxiety and NAc taurine. A ^1^H-MRS study in male mice showed that NAc taurine concentration varies with social dominance (Larrieu et al., 2017). Moreover, social competitiveness is known to be related to anxiety-like behavior in rodents (Hollis et al., 2015; Larrieu and Sandi, 2018; Van Der Kooij et al., 2018) and trait anxiety in humans (Goette et al., 2015). Moreover, highly anxious rats are less successful in dyadic competitive encounters against low anxious rats likely due to their reduced complex I and II activity and mitochondrial respiratory capacity in the NAc (Hollis et al., 2015). Importantly, taurine deficiency has been shown to impair mitochondrial complex I activity, with consequent elevation of the NADH/NAD^+^ ratio and down-regulation of energy metabolism (Schaffer et al., 2016). Importantly, we observed a negative correlation between NAc taurine and state physical fatigue. Physical fatigue is a key factor for determining success in sport competitions in humans (Knicker et al., 2011). Thus, NAc taurine might also be playing a role in human competitive performance.

Taurine can exert inhibitory neuromodulatory actions through its weak agonistic effects on GABA_A_ and glycine receptors (Bhattarai et al., 2015; Chan et al., 2014; Jia et al., 2008) [also note additional evidence of taurine’s partial agonistic actions on NMDA receptors (Mezzomo et al., 2017)]. GABAergic inhibition was shown to be controlled by taurine via its modulatory actions on the sensitivity of synaptic and extrasynaptic GABA_A_ receptors (Sergeeva et al., 2007).

The correlation between trait anxiety and NAc taurine was different to the association between the same personality trait and OL taurine. This finding suggests a regionally specific relationship between NAc taurine and trait anxiety in the human brain. Importantly, available data suggests that the striatum is particularly susceptible to low taurine concentrations (Sergeeva et al., 2007; Strolin Benedetti et al., 1991). Taurine deficiency has been shown to impair long-term plasticity in the striatum, but not in the hippocampus (Sergeeva et al., 2003). Moreover, in the rat striatum, taurine was shown to modulate dopamine uptake in a dose-dependent manner (Chen et al., 2018). Importantly, taurine supplementation can rescue GABA_A_-mediated disinhibition of striatal network activity in taurine-transporter knockout mice (Sergeeva et al., 2007). Altogether, these observations support the hypothesis that lower NAc taurine in high anxious participants might impair NAc GABAergic functioning and dopaminergic regulation.

In agreement with the literature (Sandi and Richter-Levin, 2009; Weger and Sandi, 2018), we found trait anxiety and depression to be positively associated. Importantly, we found no evidence for an association between NAc taurine and depression scores in our sample of healthy male participants. In line with previous findings (Jiang et al., 2003; Vassend et al., 2018), we observed a negative relationship between fatigue and anxiety, and we report for the first time a significant negative association between state physical fatigue and NAc taurine. State physical fatigue was assessed with the SEF 3-item scale measuring the intensity of current feelings of fatigue regarding the capacity to perform typical physical activities (O’Connor, 2006). Previous work had related taurine consumption to life stress (Sung and Chang, 2009) and physical endurance in rats (Yatabe et al., 2009) and humans (Kowsari et al., 2018). Our findings are novel in showing a negative association between taurine content in a specific brain region and physical fatigue.

Brain taurine concentrations are currently understood as a product of dietary intake, tissue specific cellular transport mechanisms and astrocytic (but possibly also neuronal) local synthesis (Pasantes-Morales, 2017). Cysteine is the main substrate for taurine synthesis. Taurine is an ingredient in various ‘energy’ drinks that typically provide 6 - 16 times the average daily amount of diet-ingested taurine (Caine and Geracioti, 2016). Most clinical trials have studied taurine effects in combination with other ingredients usually present in ‘energy’ drinks, such as caffeine. Thus, available data is insufficient to establish the specific effects of taurine consumption in healthy humans (Caine and Geracioti, 2016).

Our second main finding is the negative correlation between situational stress and NAc GABA and, consequently, a positive correlation between situational stress and E/I metabolic ratios (i.e., Glu/GABA and Glx/GABA). Our data fits with previous studies implying a major impact of stress on E/I balance, by either affecting glutamate) (see Houtepen et al., 2017, for negative findings in the PFC in a 7 T study in humans) and/or GABA (Cordero et al., 2016; Hasler et al., 2010) in different brain regions [e.g., hippocampus, prefrontal cortex (PFC), anterior cingulate (ACC)] (Popoli et al., 2012; Sandi, 2011). Particularly relevant is the reduction in GABA concentrations that has been observed following situational stress in humans reported in a ^1^H-MRS study examining PFC GABA (Hasler et al., 2010).

The observed lower NAc GABA concentrations in acutely stressed participants might be due to reduced astrocytic glutamate reuptake and consequentially lower GABA production (Goubard et al., 2011), deficiencies in enzymatic activity for GABA synthesis, and/or reduced GABA release from GABAergic afferent projections to the NAc/VS (e.g., from the ventral pallidum, amygdala, lateral hypothalamus or brainstem) (Richard et al., 2013). However, ^1^H-MRS does not allow determining the subcellular location (i.e. presynaptic nerve terminal, synaptic cleft, postsynaptic neuron or surrounding glia) of the measured metabolites. ^1^H-MRS detects tissue metabolite pools. As a speculation, the observed reduced GABA concentrations observed in our study might reflect a reduction in NAc GABAergic function resulting in reduced inhibitory potential in the NAc associated neural circuitry in humans. This interpretation is in line with the anxiolytic action of GABA receptor agonists when directly infused into the NAc/VS in rats (Lopes et al., 2012).

Although previous ^1^H-MRS studies in humans have supported opposite associations for glutamate and GABA with anxiety (Ende, 2015), our study did not identify significant associations between anxiety and glutamate, GABA or E/I balance. E/I imbalance is a key biomarker of neuropsychiatric disorders (Tatti et al., 2017). Given the strong comorbidity of stress with neuropsychiatric disorders (Peters et al., 2017; Sandi and Richter-Levin, 2009), our study suggests that self-perceived acute stress might be an important contributing factor on the E/I imbalances observed in neuropsychiatric disorders.

## Limitations

First, this study was carried out in 20-30 years-old male participants only with a modest sample size. These factors limit the interpretation of the results to the general population. Thus, the reported associations should be further studied in a female sample and in older adults. Second, our results might be relevant for the development of more effective treatments for anxiety disorders, acute stress and to reduce sensations of physical fatigue. However, we notably conducted this study in healthy individuals; whether our findings might be relevant to the clinical population remains to be established.

## Conclusions

This study shows for the first time an association between NAc taurine and GABA levels with trait anxiety and perceived acute stress in healthy individuals, suggesting the potential role of the NAc and these two metabolites in the pathophysiology of anxiety disorders. The observed negative relationship between NAc taurine and physical fatigue renders our study also relevant for models of human competitive performance.

## Supporting information

## Author contributions

A.S. and C.S. designed the research; A.S. dealt with recruitment and performed behavioral and scanner testing; L.X. developed MRS experimental conditions, performed scanners and MRS analyses; A.S., L.X., and C.S. performed statistical analyses; R.G. supervised the MRS study. A.S and C.S. wrote the paper with contributions from the other authors.

## Acknowledgements

The authors thank Olivia Zanoletti for excellent technical assistance and Dr. Fiona Hollis for her input to the experimental design.

