## Supplementary material for "Nucleus accumbens neurochemistry in human anxiety: A 7 T ^1^H-MRS study"

**SUPPLEMENTARY METHODS AND MATERIALS**

### Psychometric questionnaires

State and trait anxiety were assessed with the State-Trait Anxiety Inventory (STAI) (Spielberger, 2010) after arriving in the lab, 90 min before ^1^H-MRS. Self-perceived stress was measured with a 10-point rating scale (“How stressed do you feel at the moment?” ranging from “not at all” to “very much”) on the scanner bed in a seated position via written self-report, 30 min before ^1^H-MRS. Social anxiety and depression were assessed 60 min after ^1^H-MRS with the Liebowitz Social Anxiety Scale (LSAS) (Heimberg et al., 1999). At this time point, participants also filled responses for the Beck Depression Inventory (BDI) (Beck, 1961) and the Mental and Physical State Energy and Fatigue Scales (SEF) (Loy and O’Connor, 2016). State anxiety and acute stress were measured twice; at the NAc ^1^H-MRS session and at the OL ^1^H-MRS session. Tables S1 and S2 show the phenotypic characteristics of participants for the questionnaires filled in each of the two experimental sessions.

**Salivary cortisol**

Saliva was collected three times (baseline, pre and post MRS acquisition) on the first experimental day. Participants were instructed to place the saliva cotton swab (Salimetrics, State College, PA, USA) medially under the tongue for 2 min. Samples were immediately stored on dry ice and then frozen at -70°C. Free cortisol concentrations were determined with the Salimetrics Salivary Cortisol Enzyme Immunoassay Kit. Area under the curve values (AUC_G_, AUC_I_) were computed according to Pruessner et al. (Pruessner et al., 2003). AUC_G_ denotes the area under the curve with respect to the baseline value, and AUC_I_ denotes the area under the curve with respect to increase (5). Table S3 includes the descriptive statistics for the salivary cortisol concentrations.

### 1H magnetic resonance spectroscopy (1H-MRS)

^1^H-MRS was performed on a 7 T/68 cm MR scanner (Magnetom, Siemens Medical Solutions, Erlangen, Germany) equipped with head gradients with maximum strengths of 50 mT/m and 2^nd^-order shims of up to 13.5 mT/m^2^ for Z^2^ and 11.8 mT/m^2^ for ZX, ZY, X^2^-Y^2^ and XY shims. A single-channel quadrature transmit and a 32-channel receive coil (Nova Medical Inc., MA, USA) were used for ^1^H-MRS in the NAc. A home-built ^1^H quadrature transmit/receive surface coil was used for OL ^1^H-MRS. T_1_-weighted anatomical images were acquired with a magnetization-prepared rapid gradient-echo (MP2RAGE) sequence [repetition time (TR) = 5500 ms, echo time (TE) = 1.87 ms, inversion time (TI)_1_ = 750 ms, TI_2_ = 2350 ms, α_1_ = 4°, α_2_ = 5°, 1 × 1 × 1 mm resolution, matrix size = 210 × 210 × 160, acquisition time (TA) = 7 min 20 sec] to position the volume-of-interest (VOI) and for subsequent VOI tissue segmentation (Marques et al., 2010).

To ensure the full inclusion of the NAc, the VOI was defined by the third ventricle medially, the subcallosal area inferiorly, and the body of the caudate nucleus and the putamen laterally and superiorly, in line with definitions of NAc anatomy identifiable on MRIs (Neto et al., 2008) (Figure 1A).

B_0_ field inhomogeneities within the VOI were minimized using 1^st^- and 2^nd^-order shims with the fast, automatic shim technique using echo-planar signal readout for mapping along projections FAST(EST)MAP sequence (Gruetter, 1993; Gruetter and Tkáč, 2000). Higher 2^nd^-order shim values were required for the NAc relative to the OL, in particular for the XY, ZY, and ZX shims (Figure S1).

Transmit reference voltage was then calibrated to obtain the maximal water signal from the VOI. Then, outer volume suppression (OVS) bands were placed around the VOI to reduce spectral contaminations from lipid and water signals from the VOI periphery. The variable pulse power and optimized relaxation delays (VAPOR) sequence was applied for water signal suppression (Tkáč and Gruetter, 2005). ^1^H-MR spectra were acquired with the semi-adiabatic spin-echo full-intensity acquired localized (semi-adiabatic SPECIAL) sequence (Xin et al., 2013). In the NAc (VOI = 14 × 10 × 13 mm^3^, TR/TE = 6500/16 ms, bandwidth = 4000 Hz, vector size = 2048 pts, average of 8, 32 blocks) and in the OL (VOI = 25 × 20 × 20 mm^3^, TR/TE = 8000/16 ms, bandwidth = 4000 Hz, vector size = 2048 pts, average of 2, 32 blocks). The unsuppressed water signal was acquired as an internal reference for the metabolite quantification and eddy current correction in both brain regions.

Localized single-voxel ^1^H-MR spectra from the NAc were obtained from twenty-seven participants. ^1^H-MRS in the OL was performed as the experimental control (*N* = 17).

**SUPPLEMENTARY FIGURES**

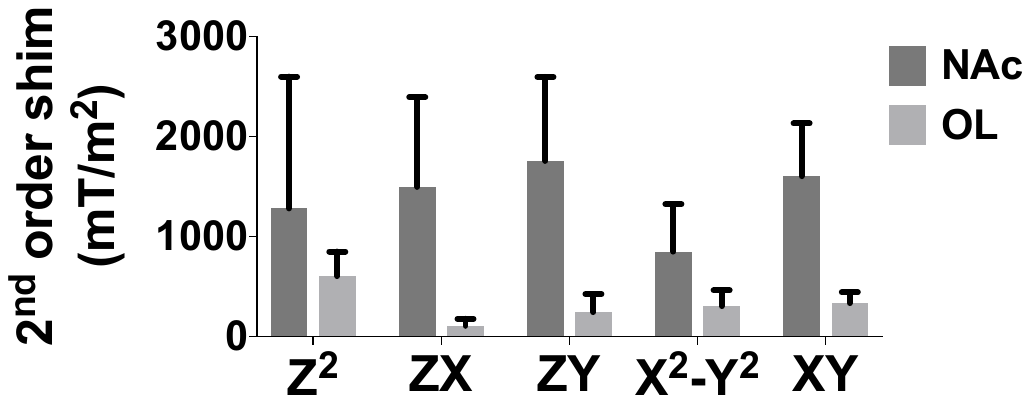

**Figure S1. Second order shim strength for the nucleus accumbens and the occipital lobe.** Means and standard deviations (SDs) are shown. NAc, nucleus accumbens, *N* = 27; OL, occipital lobe, *N* = 17.

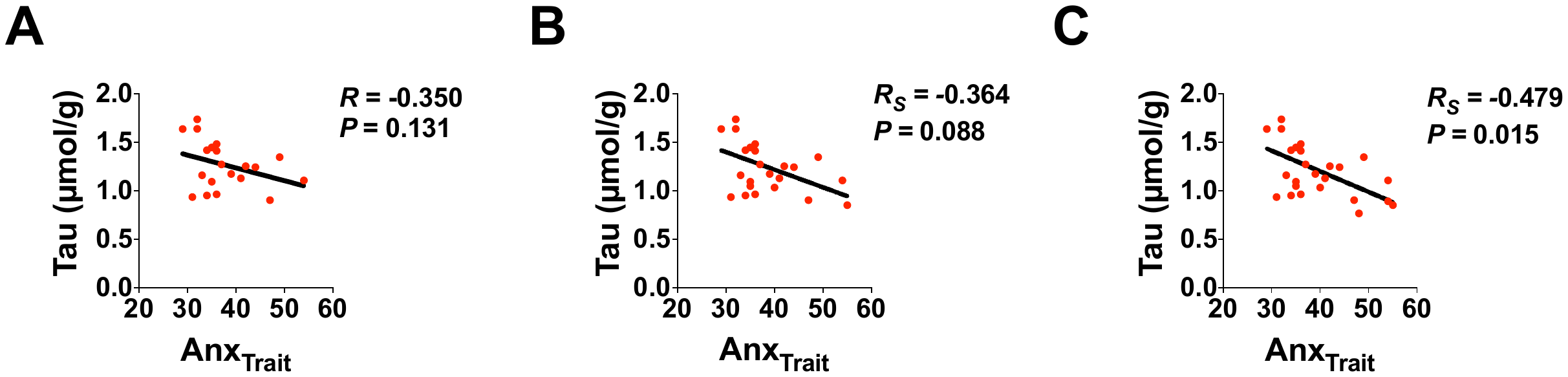

**Figure S2. Associations of trait anxiety with nucleus accumbens taurine.** Scatter plots of the associations of trait anxiety with taurine (
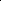
 20 % CRLB), *N* = 20 (**A**); taurine (
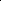
 30 % CRLB), *N* = 23 (**B**); taurine (
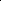
 50% CRLB), *N* = 25 (**C**). Anx_Trait_; trait anxiety (STAI-T). Tau, taurine. *R*, Pearson’s correlation coefficient; *R_S_*, Spearman’s correlation coefficient. *P*-values are uncorrected.

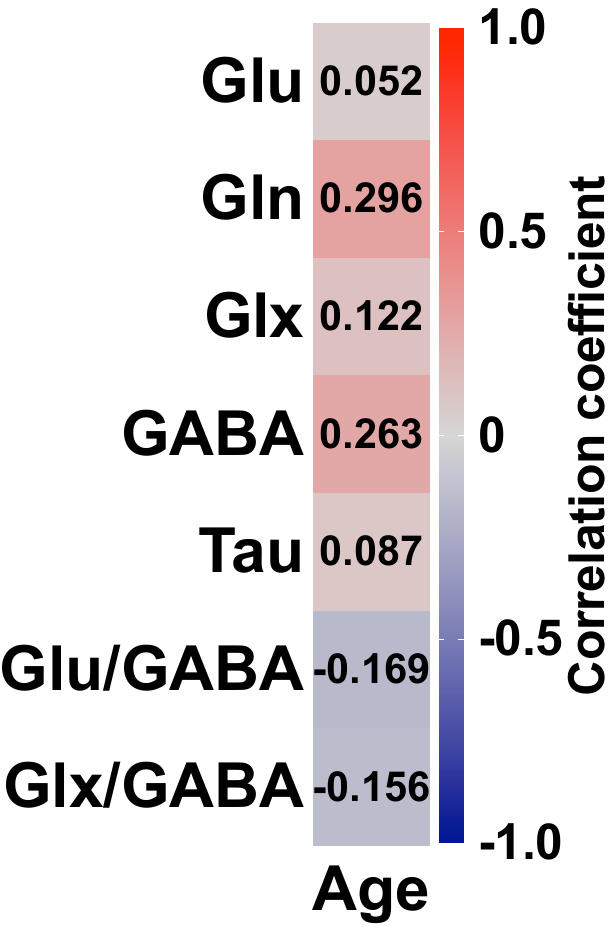

**Figure S3. Lack of significant associations between participants’ age (20-30 years-old) and nucleus accumbens metabolites.** Glu, glutamate; Gln, glutamine; Glx, Glu+Gln; GABA, gamma-aminobutyric acid; Tau, taurine. Each cell contains Spearman’s correlation coefficient corresponding to the colour scale.

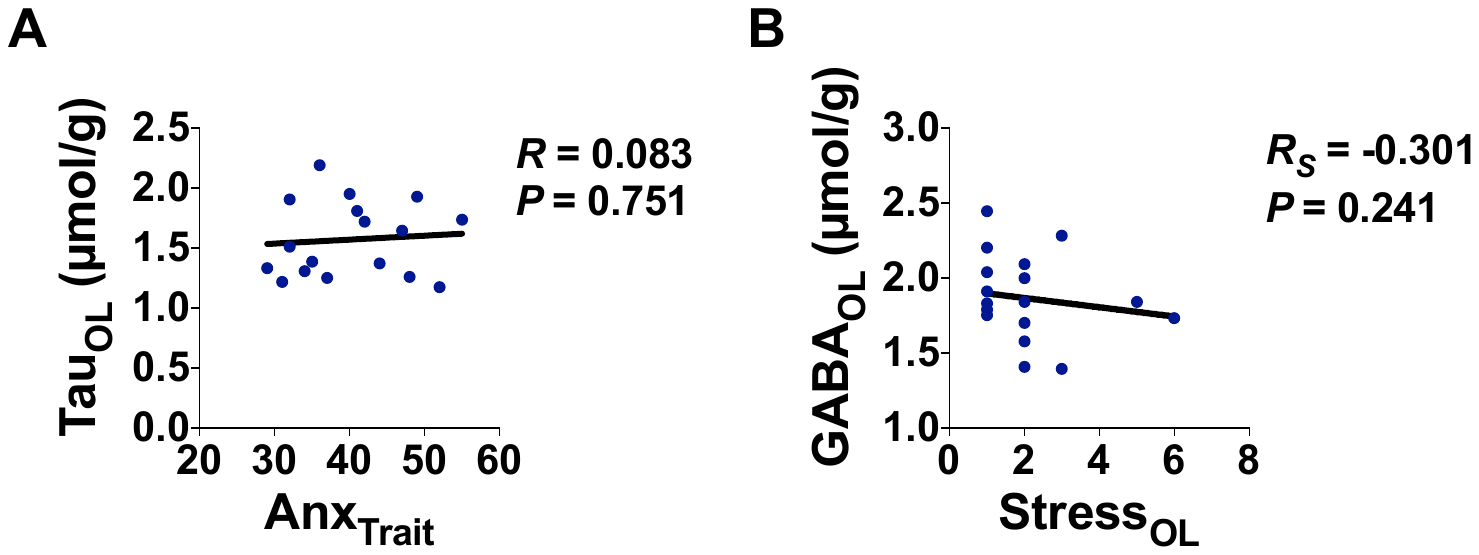

**Figure S4. Lack of correlation between anxiety trait and taurine and between stress and GABA in the occipital lobe.** (**A**) Scatter plot of trait anxiety and taurine. (**B**) Scatter plot of stress and GABA. Anx_Trait_, trait anxiety (STAI-T); Stress, self-perceived acute stress. Tau, taurine; GABA, gamma-aminobutyric acid. OL, occipital lobe. *N* = 17. *R*, Pearson’s correlation coefficient; *R_S_*, Spearman’s correlation coefficient. **P* < 0.05; uncorrected.

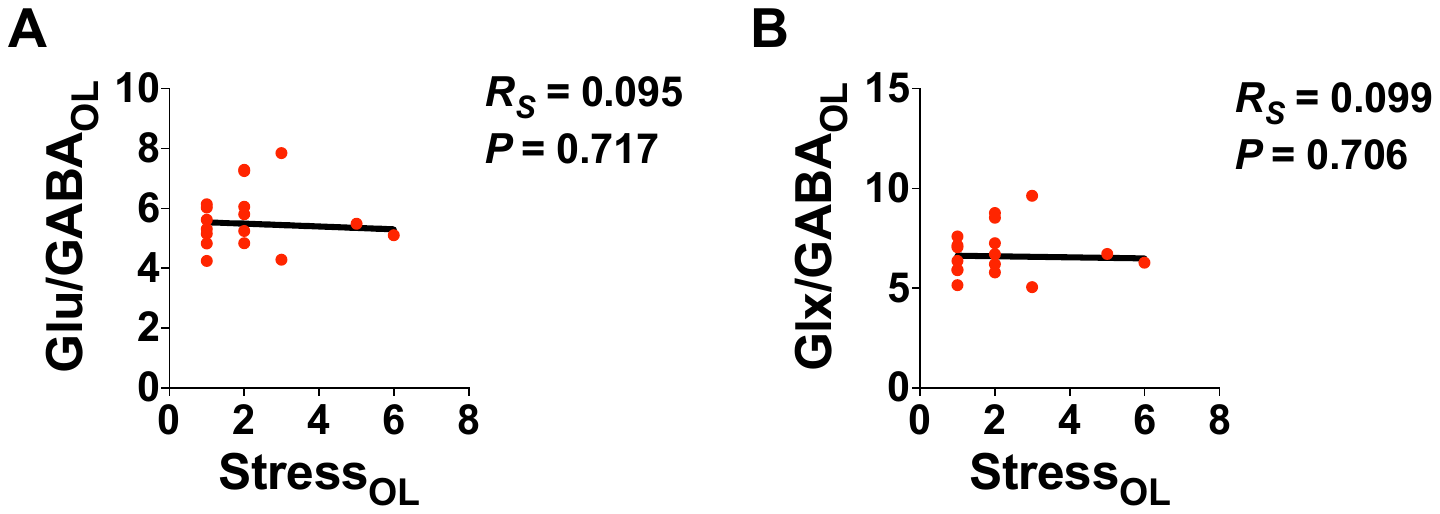

**Figure S5. Lack of significant correlation between stress and excitation/inhibition balance in the occipital lobe.** (**A**) Scatter plot of stress and Glu/GABA. (**B**) Scatter plot of stress and Glx/GABA. Stress, self-perceived acute stress. Glu, glutamate; Gln, glutamine; Glx, Glu+Gln; GABA, gamma-aminobutyric acid. OL, occipital lobe. *N* = 17. *R_S_*, Spearman’s correlation coefficient.

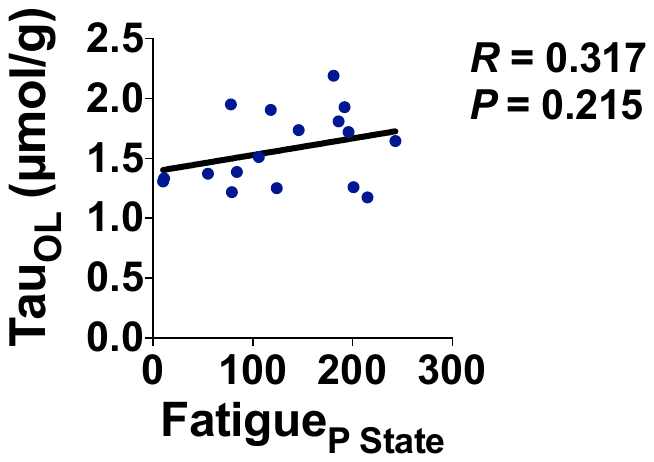

**Figure S6. Associations between state physical fatigue and occipital lobe taurine.** (**A**) Scatter plot of state physical fatigue and taurine. Fatigue_P State_, state physical fatigue; Tau, taurine. OL, occipital lobe. *N* = 17. *R*, Pearson’s correlation coefficient.

**SUPPLEMENTARY TABLES**

**Table S1. Phenotypic characteristics. Anx_State_, state anxiety (STAI-S); Anx_Trait_, trait anxiety (STAI-T); Anx_Social_, social anxiety (LSAS); Stress, self-perceived acute stress; Energy_M State_, state mental energy; Energy_M Trait_, trait mental energy; Energy_P State_, state physical energy; Energy_P Trait_, trait physical energy; Fatigue_M State_, state mental fatigue; Fatigue_M Trait_, trait mental fatigue; Fatigue_P State_, state physical fatigue; Fatigue_P Trait_, trait physical fatigue. *M*, mean; *SD,* standard deviation.**

|  | ***N*** | ***Min*** | ***Max*** | ***M*** | ***SD*** |
| --- | --- | --- | --- | --- | --- |
| ***Age*** | 38 | 20 | 30 | 23.03 | 2.93 |
| ***Anx_State_*** | 38 | 20 | 50 | 30.26 | 6.98 |
| ***Anx_Trait_*** | 38 | 28 | 56 | 40.21 | 7.86 |
| ***Anx_Social_*** | 34 | 18 | 89 | 47.68 | 17.40 |
| ***Stress*** | 35 | 1 | 7 | 3.26 | 1.69 |
| ***Depression*** | 35 | 0 | 37 | 9.71 | 7.28 |
| ***Energy_M State_*** | 35 | 66 | 242 | 158.71 | 50.35 |
| ***Energy_M Trait_*** | 35 | 4 | 12 | 7.69 | 2.05 |
| ***Energy_P State_*** | 35 | 70 | 211 | 160.03 | 40.92 |
| ***Energy_P Trait_*** | 35 | 5 | 12 | 8.17 | 1.65 |
| ***Fatigue_M State_*** | 35 | 16 | 222 | 119.54 | 61.65 |
| ***Fatigue_M Trait_*** | 35 | 0 | 9 | 4.20 | 2.08 |
| ***Fatigue_P State_*** | 35 | 10 | 243 | 124.89 | 66.52 |
| ***Fatigue_P Trait_*** | 35 | 1 | 6 | 3.54 | 1.62 |

**Table S2. State anxiety and perceived stress at the ^1^H-MRS session in the occipital lobe. Anx_State_, state anxiety (STAI-S); Stress, self-perceived acute stress. *M*, mean; *SD,* standard deviation.**

|  | ***N*** | ***Min*** | ***Max*** | ***M*** | ***SD*** |
| --- | --- | --- | --- | --- | --- |
| ***Anx_State_*** | 18 | 20 | 42 | 30 | 7.41 |
| ***Stress*** | 18 | 1 | 6 | 2.06 | 1.43 |

**Table S3. Salivary cortisol concentrations and changes over time. Cort_BL_, cortisol baseline; Cort_AUC-G_, cortisol area under the curve with respect to ground; Cort_AUC-I_, cortisol area under the curve with respect to increase. *M*, mean; *SD,* standard deviation.**

|  | ***N*** | ***Min*** | ***Max*** | ***M*** | ***SD*** |
| --- | --- | --- | --- | --- | --- |
| ***Cort_BL_*** | 28 | 0.06 | 0.62 | 0.24 | 0.14 |
| ***Cort_AUC-G_*** | 22 | 8.33 | 43.29 | 21.90 | 11.25 |
| ***Cort_AUC-I_*** | 22 | -20.19 | 13.03 | -2.83 | 7.66 |

**Table S4. Nucleus accumbens metabolite concentrations.** Absolute concentrations and CRLBs are given as mean ± standard deviation. Glu, glutamate; Gln, glutamine; GABA, gamma-aminobutyric acid; Tau, taurine. CRLB, Cramér-Rao lower bound. *N* = 27. Metabolite concentrations and absolute CRLBs are given in μmol/g. *M*, mean.

|  | ***N*** | ***Min*** | ***Max*** | ***M*** | ***Absolute CRLB*** | ***CRLB (%)*** |
| --- | --- | --- | --- | --- | --- | --- |
| ***Glu*** | 27 | 7.67 | 11.63 | 9.79 ± 1.09 | 0.23 ± 0.06 | 2.41 ± 0.57 |
| ***Gln*** | 27 | 2.64 | 6.82 | 4.45 ± 1.05 | 0.21 ± 0.04 | 4.89 ± 0.89 |
| ***GABA*** | 27 | 1.09 | 3.17 | 2.22 ± 0.53 | 0.24 ± 0.04 | 11.41 ± 2.74 |
| ***Tau*** | 27 | 0.32 | 1.74 | 1.14 ± 0.33 | 0.21 ± 0.04 | 21.52 ± 14.91 |

**Table S5. Correlation matrix of nucleus accumbens metabolites. Glu, glutamate; Gln, glutamine; Glx, Glu+Gln; GABA, gamma-aminobutyric acid; Tau, taurine. Pearson’s correlation coefficients are indicated. *N* = 27**

|  | ***Glu*** | ***Gln*** | ***Glx*** | ***GABA*** | ***Tau*** |
| --- | --- | --- | --- | --- | --- |
| ***Glu*** | 1 | 0.802**** | 0.951**** | 0.604*** | 0.144 |
| ***Gln*** | 0.802**** | 1 | 0.947**** | 0.581 | -0.049 |
| ***Glx*** | 0.951**** | 0.947**** | 1 | 0.624*** | 0.052 |
| ***GABA*** | 0.604*** | 0.581** | 0.624*** | 1 | 0.210 |
| ***Tau*** | 0.144 | -0.049 | 0.052 | 0.210 | 1 |

**P* < 0.05; ***P* < 0.01; ****P* < 0.001; *****P* < 0.0001.

**Table S6. Occipital lobe metabolite concentrations and E/I balance.** Glu, glutamate; Gln, glutamine; GABA, gamma-aminobutyric acid; Tau, taurine. Metabolite concentrations and absolute CRLBs are given in μmol/g. *M*, mean.

|  | ***N*** | ***Min*** | ***Max*** | ***M*** | ***Absolute CRLB*** | ***CRLB (%)*** |
| --- | --- | --- | --- | --- | --- | --- |
| ***Glu*** | 17 | 8.85 | 12.40 | 10.40 ± 0.86 | 0.14 ± 0.05 | 1.44 ± 0.51 |
| ***Gln*** | 17 | 1.39 | 2.91 | 2.11 ± 0.41 | 0.12 ± 0.02 | 6.06 ± 1.39 |
| ***GABA*** | 17 | 1.39 | 2.45 | 1.87 ± 0.29 | 0.13 ± 0.02 | 7.38 ± 1.31 |
| ***Tau*** | 17 | 1.17 | 2.19 | 1.57 ± 0.31 | 0.11 ± 0.02 | 7.25 ± 1.73 |

**Table S7. Correlation matrix of occipital lobe metabolites.** Glu, glutamate; Gln, glutamine; Glx, Glu+Gln; GABA, gamma-aminobutyric acid; Tau, taurine. Pearson’s correlation coefficients are indicated. *N* = 17.

**P* < 0.05; ***P* < 0.01; ****P* < 0.001; *****P* < 0.0001.

|  | ***Glu*** | ***Gln*** | ***Glx*** | ***GABA*** | ***Tau*** |
| --- | --- | --- | --- | --- | --- |
| ***Glu*** | 1 | 0.623** | 0.961**** | -0.010 | 0.337 |
| ***Gln*** | 0.623** | 1 | 0.814**** | 0.060 | 0.363 |
| ***Glx*** | 0.961**** | 0.814**** | 1 | 0.014 | 0.378 |
| ***GABA*** | -0.010 | 0.060 | 0.014 | 1 | 0.094 |
| ***Tau*** | 0.337 | 0.363 | 0.378 | 0.094 | 1 |
